# Uncovering Conceptual Biases in DNA Stabilization: A Student-Led Investigation

**DOI:** 10.64898/2026.01.15.699716

**Authors:** Charlotte Polo, Ameeta Thandi, Olivia Chandler, Paula Lugert, Alyssa Hammoud, Theertha Madhi, Malena Ayala, A.J. Berrigan, Andrew A.Y. Chen, Kate Gillett, Sohan Sanjeev, Mya Sareen, Sean Yu, Yang-yang Zuo, Shawn Xiong

## Abstract

Deoxyribonucleic acid (DNA) stands as one of the most foundational concepts in life sciences, essential for students to master. However, when surveyed about the forces that stabilize the double-stranded DNA structure, many students exhibited a conceptual bias—favoring base pairing as the primary stabilizing force, while overlooking the equally critical role of base stacking interactions. To investigate the origins of this misconception, students conducted an analysis of 35 widely used textbooks. Their findings revealed that one-third of these texts explicitly emphasized base pairing as the sole stabilizing force in their written content. Furthermore, two-thirds of the textbook contained illustrations that reinforced this bias, visually highlighting base pairing while neglecting base stacking. Recognizing this bias, students conducted literature research to gain a more accurate and nuanced understanding of DNA stabilization. Through their research, students identified three concept areas—DNA structure and function, environmental effects on DNA, and DNA-protein interactions—to illustrate how base pairing and base stacking work in concert to stabilize the antiparallel double helical structure of DNA. This interplay between base pairing and base stacking is crucial not only for the structural integrity of DNA, but also for its biological functionality. By addressing this conceptual bias, we aim to promote a more balanced and scientifically accurate representation of DNA stabilization in educational materials.

## Introduction

Double stranded deoxyribonucleic acid (dsDNA) is a fundamental biomolecule, providing the basis for life on earth. DNA’s function is intricately linked to its structure, which is stabilized by covalent and non-covalent interactions. In 1953, James Watson and Francis Crick published a proposal for DNA structure based on X-ray diffraction data collected by Rosalind Franklin, now known as the double helix model (Watson & Crick, 1953; Franklin & Gosling, 1953; Figure 1). Their proposed structure consists of two antiparallel sugar-phosphate chains connected by phosphodiester linkages; both intertwined in right-handed helices. Backbone sugars are bound to heterocyclic nucleobases, composed of single and double-ringed carbon backbones (pyrimidines and purines, respectively), via an N-glycosidic linkage. These nucleobases are oriented parallel, pairing selectively such that purines bind pyrimidines, allowing for the transmission of genetic information. In canonical base pairing, adenine pairs with thymine (AT) through two hydrogen bonds (H-bonds), while guanine pairs with cytosine (GC) through three H-bonds. The double helix is stabilized through the hydrophobic effect, whereby the hydrophilic phosphate groups interact with the polar solvent, and the hydrophobic nucleobase are buried.

**Figure 1.**
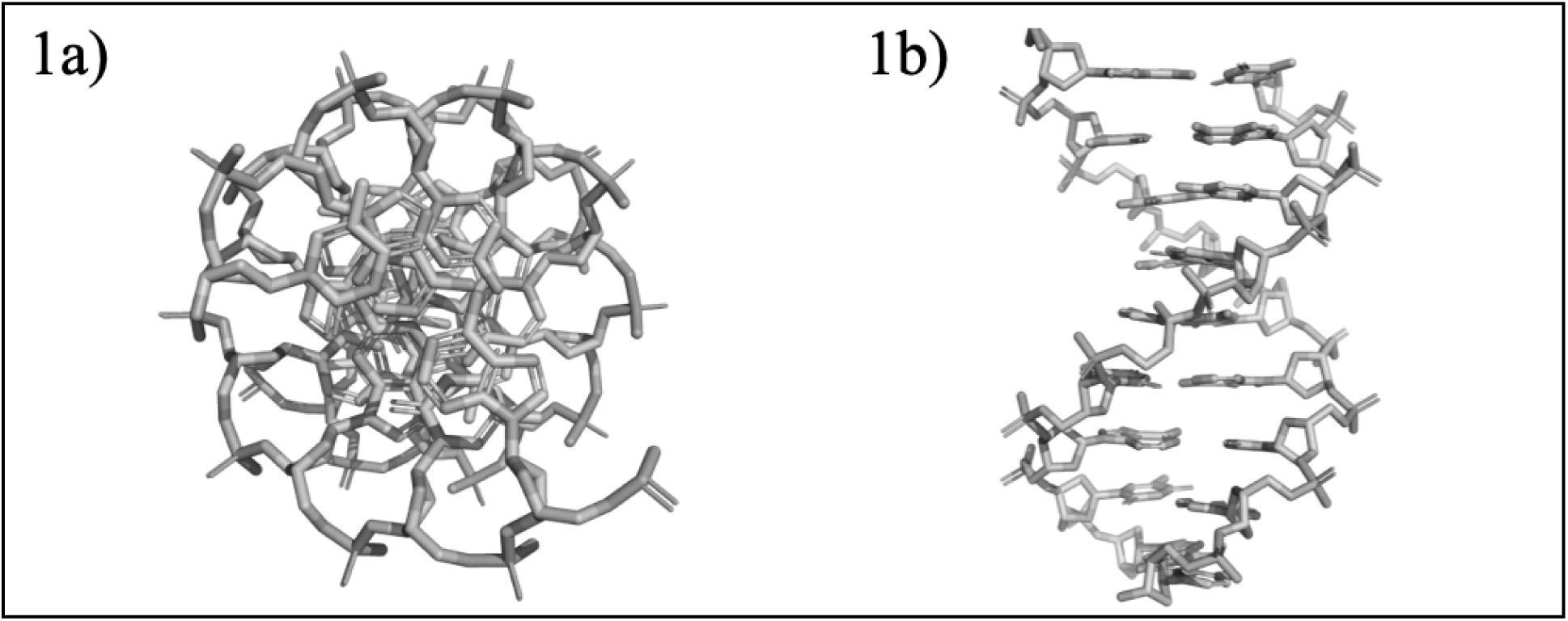
Molecular models of double helix DNA. a) Bird’s-eye and b) side-on views of the Watson-Crick model for double helix DNA, generated from a random sequence using the “fnab” command in PyMOL.

DNA utilizes four distinct stabilizing forces. For instance, the phosphate groups within the backbone of the helix have been found to establish stable electrostatic interactions with common intracellular cations such as sodium and magnesium when deprotonated at physiological condition (Huguet et al., 2017; Feughelman et al., 1955). Additionally, these phosphate groups form phosphodiester linkages with deoxyribose sugars through condensation reactions, providing stability to the structure through strong covalent bonds (Elliott & Elliott, 2005). Nucleobases influence the stability of dsDNA through base pairing (facilitated by H-bonds) and base stacking (facilitated by orbital overlap and van der Waals forces between stacked bases) (Krueger et al., 2006; Poater et al., 2014; Li et al., 2009). While the four forces mentioned above (phosphodiester bonds, ionic interactions, base pairing, and base stacking) work synergistically to stabilize DNA, the amount each force contributes to total stability remains a subject of ongoing debate and investigation.

Base pairing, mediated by H-bonds between complementary bases, AT and GC, reveals its critical role in DNA replication, transcription, and in the fidelity of genetic information flow (Petersheim & Turner, 1983). The H-bonds between base pairs help reinforce the DNA strands, particularly at the ends, where base stacking interactions are weaker (Matray & Kool, 1998). This function is essential for maintaining the structural integrity of linear DNA by reducing end fraying and strand separation (Matray & Kool, 1998). In contrast, base stacking interactions, driven by van der Waals forces and the hydrophobic effect, contribute significantly to the enthalpic stability of dsDNA (Krueger et al., 2006). These stacking interactions are enhanced by the overlap of π-orbitals between neighboring bases and the additive contributions of van der Waals forces, providing a strong stabilizing effect, and thus crucial for the structural integrity and overall stability of the DNA duplex (Friedman & Honig, 1992).

Student perspectives on the stabilizing forces of dsDNA offer another lens to assess the broader impact of the base pairing versus base stacking discourse. Survey data collected from biochemistry and molecular biology undergraduates at an Atlantic Canadian university revealed that many second-year students identified base pairing as the primary stabilizing force (Figure 2a). A smaller proportion recognized both base pairing and base stacking as contributing to stability, while very few considered base stacking alone to be significant. Notably, this trend persisted into their third year of study, with little to no improvement in their understanding (Figure 2b). Despite increased academic exposure, students continued to place disproportionate emphasis on base pairing; these results prompted our inquiry into the following research questions:

1. How might textbook representations both in writing and in figure contribute to students’ oversimplified view on DNA stability?
2. How does the current literature represent the intermolecular forces involved in DNA stability?

**Figure. 2.**
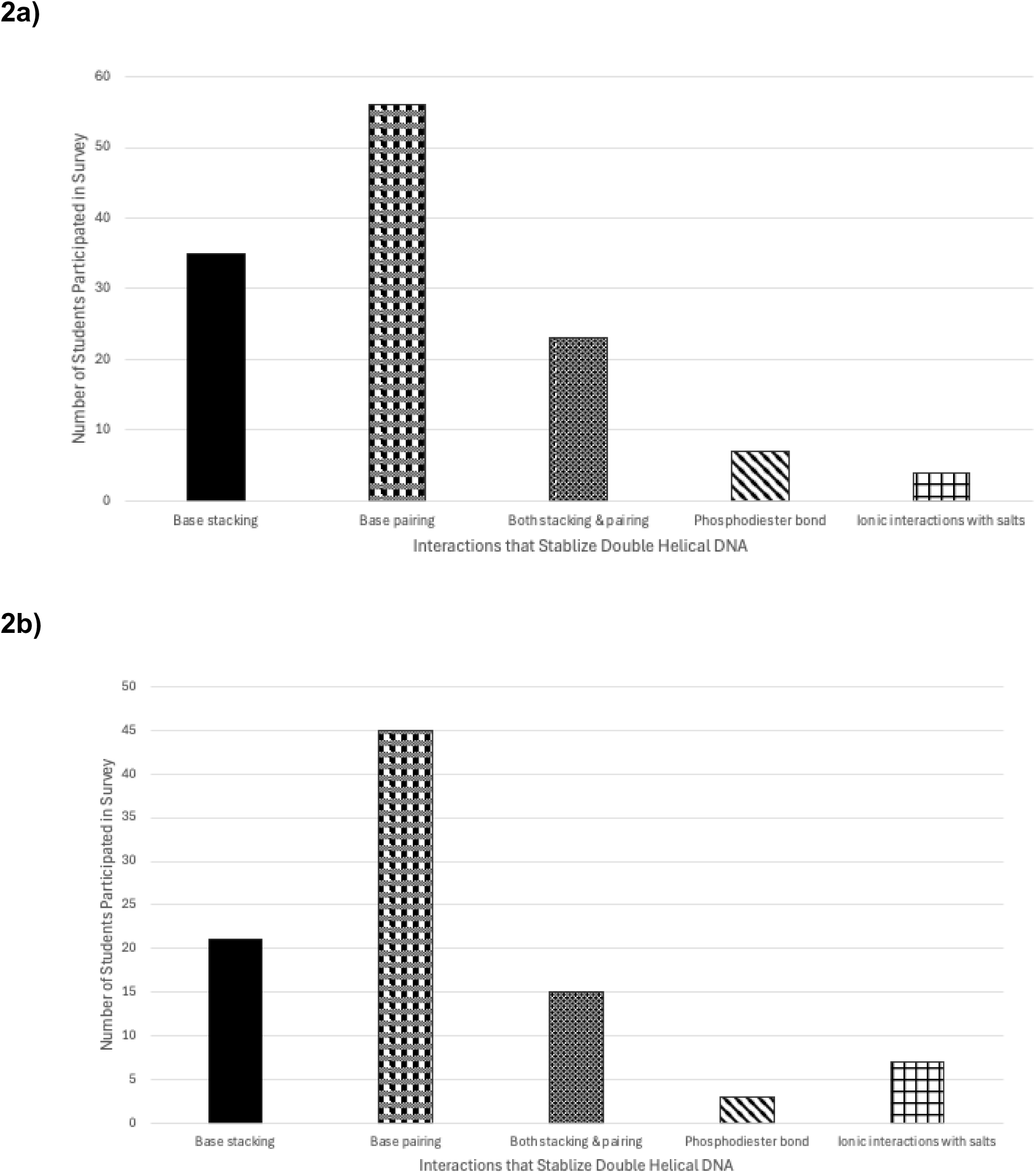
Students’ perception of the intermolecular forces that stabilize double stranded DNA. a) Survey of an Atlantic Canadian university biochemistry and molecular biology undergraduate students in a second-year biochemistry course regarding what they believed to be the stabilizing force(s) of double stranded DNA. b) Survey of the same cohort of undergraduate students in their third year on what they believed to be the stabilizing force(s) of double stranded DNA. X-axis shows the categorization of different biases, and Y-axis shows the number of students participated in the survey.

This study investigates textbooks and literature to answer these two research questions. It was hypothesized that textbook representations of DNA in writing and/or in figures may have contributed to students’ persistent and oversimplified views on DNA structural stability, biasing towards base pairing as the dominant or sometimes only stabilizing force in DNA. In the first part of this paper, 35 textbooks were examined, considering how the texts and figures influence students’ perceptions on the stability of DNA. Subsequently, the second half of the paper presents three key areas that exemplify a holistic depiction of DNA stability—DNA structure and function, environmental effects on DNA, and DNA-protein interaction— with the aim to address the gap in students’ understanding and prevent the perpetuation of conceptual bias.

## Methods

### Survey of student perception on DNA stability

Survey of student perception on DNA stability was conducted in a second year, required biochemistry laboratory course (n=125) as part of the informal pre-lab prior knowledge check-in and repeated with the same cohort in their third year required biochemistry laboratory course (n=91). Students in the labs were presented with the following question on the screen:

Question: Which one of the following interactions is the most important in stabilizing double stranded DNA?

A. Base stacking
B. Base pairing
C. Both base stacking and base pairing
D. Phosphodiester bond
E. Ionic Interaction

Students were encouraged to submit their choice voluntarily and anonymously by scanning a QR code on the screen through Microsoft Forms.

### Description of the Research Program

Student researchers were recruited voluntarily through alternative assessment as part of a year-long laboratory experience at a third-year level. As part of the alternative assessment group, students used their research and literature review to replace the credits that would be normally evaluated through writing lab reports. Since both the alternative and regular assessments were aimed to help students develop scientific writing skills, students from either group were unlikely to be disadvantaged throughout the laboratory learning process. During literature research and review, students focused on gaining a holistic and unbiased view on the forces that stabilize dsDNA from the current literature.

In total, 14 students chose to pursue the alternative assessment guided by a series of scaffolded formative assessments including bibliography, article summary, argument development, multiple drafts at different stages, and eventually the final report, which was used as the basis for developing the current manuscript (Hati & Bhattacharyya, 2024). Students were encouraged to work in pairs at the early stages of research and article summary and then were promoted to work in slightly larger groups, up to 6 people, at the later stage upon incorporating community data contributed by all 14 members. The scaffolded formative assessments were aimed to keep students on track throughout the year-long project while receiving timely feedback informing the next stage of writing. Two students were recruited over the summer of 2025 in a special topics course to take charge on compiling and finalizing the manuscript. The compiled document was read, edited, and revised by all students repeatedly, and the final editing was done by the instructor before submitting the manuscript for peer review.

### Development of the decision tree to triage textbooks

The first round of textbook analysis was carried out in student groups as part of the scaffolded writing assignment as describe before. In the scaffolded writing assignments, students were asked specifically to examine the textbooks available to them and devise their own criteria for categorizing textbooks into either biased towards base pairing or biased towards base stacking. As part of the feedback to students’ writing assignments, the instructor pooled categorization criteria from different groups and made them available to all students in the alternative assessment group. Students were then asked to use the pooled categorization criteria to reassess their textbooks during their final revision. This did not yield a consistent categorization of textbooks among different student groups. To resolve this problem, three coders (two students and one instructor) were recruited over the summer to re-examine students’ categorization criteria and the cause of inconsistency. It was found the cause of the inconsistency in categorization was due to one third of the available textbooks having a more balanced representation of base pairing vs. base stacking, leading to ambivalent categorization. As a result, the coders included a third category, “unbiased”. Three coders further cleaned up the categorization criteria by deleting redundant criteria and by restricting the analysis to a specific chapter of the textbooks. As a result, a series of Yes and No question criteria were constructed into a decision tree for textbook analysis, see Figure 3.

**Figure 3.**
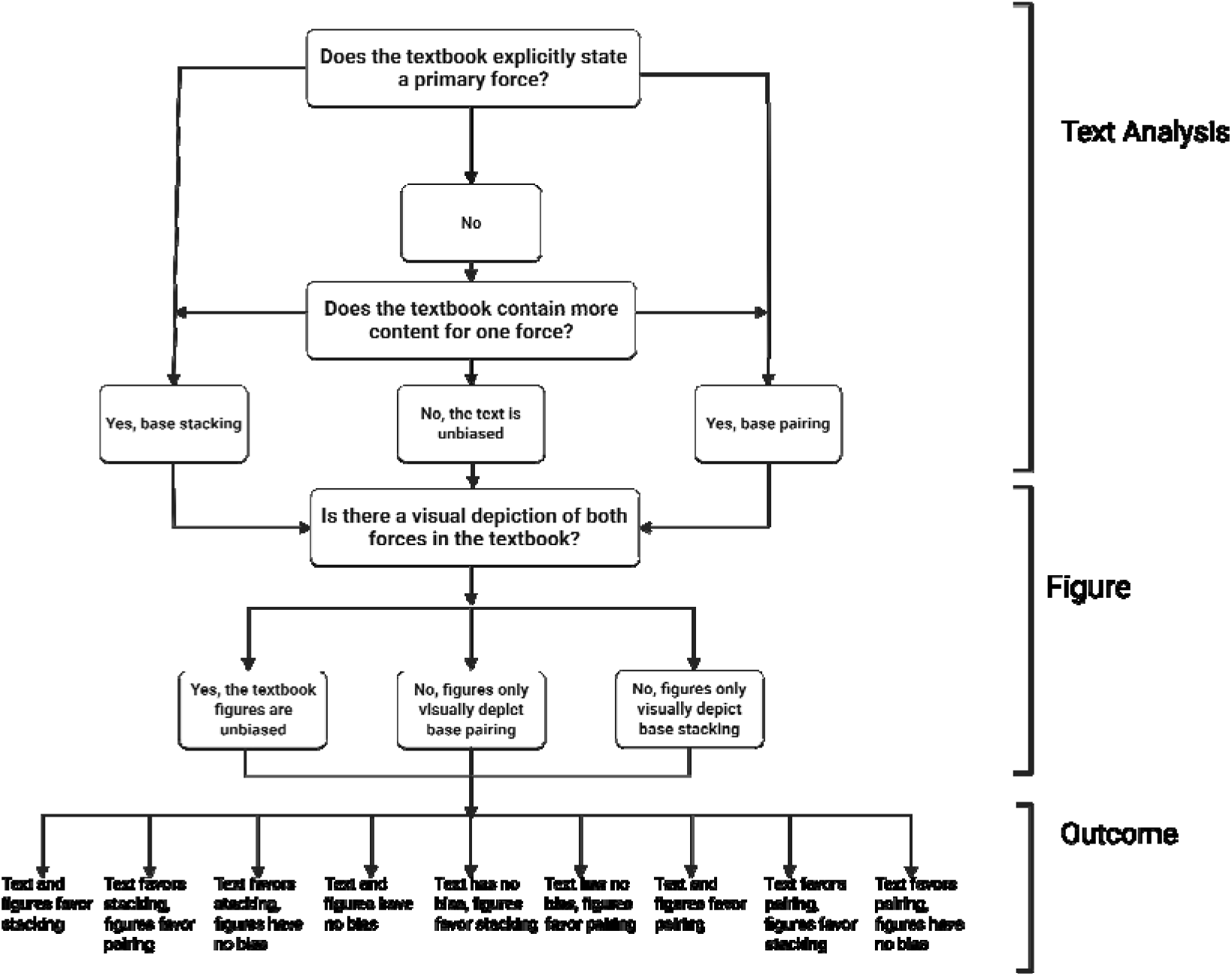
Decision tree used to assess base pairing or base stacking biases in textbooks.

### Textbook Analysis

The textbooks selected for analysis spanned a range of disciplines within the natural sciences, including biochemistry, molecular biology, genetics, biology, and biological chemistry. (Ahern et al., 2018; Alberts et al., 2022; Anthony-Cahill et al., 2019; Armstrong, 1983; Berg et al., 2002; Berg et al., 2023; Boyer, 1999; Campbell & Farrell, 2003; Campbell & Reece, 2002; Cox et al., 2025; Cox et al., 2015; Devlin, 2002; Dow et al., 1996; Elliott & Elliott, 1997; Elliott & Elliott, 2005; Ferrier, 2014; Garrett & Grisham, 2005; Horton et al., 2012; Jakubowski & Flatt, 2018; Klostermeier & Rudolph, 2018; McKee & McKee, 2016; McMurry et al., 2013; McMurry et al., 2017; Miesfeld & McEvoy, 2017; Miesfeld & McEvoy, 2021; Nelson & Cox, 2008; Nelson & Cox, 2021; Panini, 2013; Pierce, 2019; Pratt & Cornely, 2021; Tansey, 2020; Timberlake, 2010; Tymoczko et al., 2019; Voet & Voet, 2011; Voet et al., 2016). The objective of the analysis was to identify potential biases in how base stacking and base pairing interactions are presented as stabilizing forces in dsDNA. Subsequent text and figure analysis considered only textbook chapters devoted to DNA structure. To assess textual bias, the aforementioned decision tree was employed. Textbooks were categorized based on whether they presented an explicit preference for base stacking or base pairing as the primary stabilizing interaction. If the text explicitly stated that one interaction contributed more significantly to DNA stability than the other, it was classified as biased towards that force.

In cases where no such explicit statement was provided, the relative volume of discussion dedicated to each interaction was evaluated. Textbooks that devoted substantially more content to one force were considered biased towards that force. If there was no explicit statement, and the volumes of discussion for both forces were equivalent, textbook content was classified as unbiased.

In the second part of the analysis, figures depicting dsDNA were evaluated for visual bias. A figure was considered biased towards base pairing, if it only highlighted the interactions between two parallel nucleobases, such as using dashed lines to represent H-bonds (Figure 4a). On the other hand, a figure was categorized as biased towards base stacking, if it only represented the interactions between the co-planar nucleobases vertically, such as using lines, shading, electron clouds or labels to highlight π-π interactions within van der Waals’ distance (Figure 4b). A figure was deemed unbiased if it clearly illustrated both base pairing and base stacking, using visual indicators that allow students to recognize its presence and significance without prior knowledge of the stabilizing forces of DNA (Figure 4c). Figure captions were excluded from this analysis, as it was found that there was a poor correlation between the mention of base stacking in the figure caption and its representation within the figure itself.

**Figure 4.**
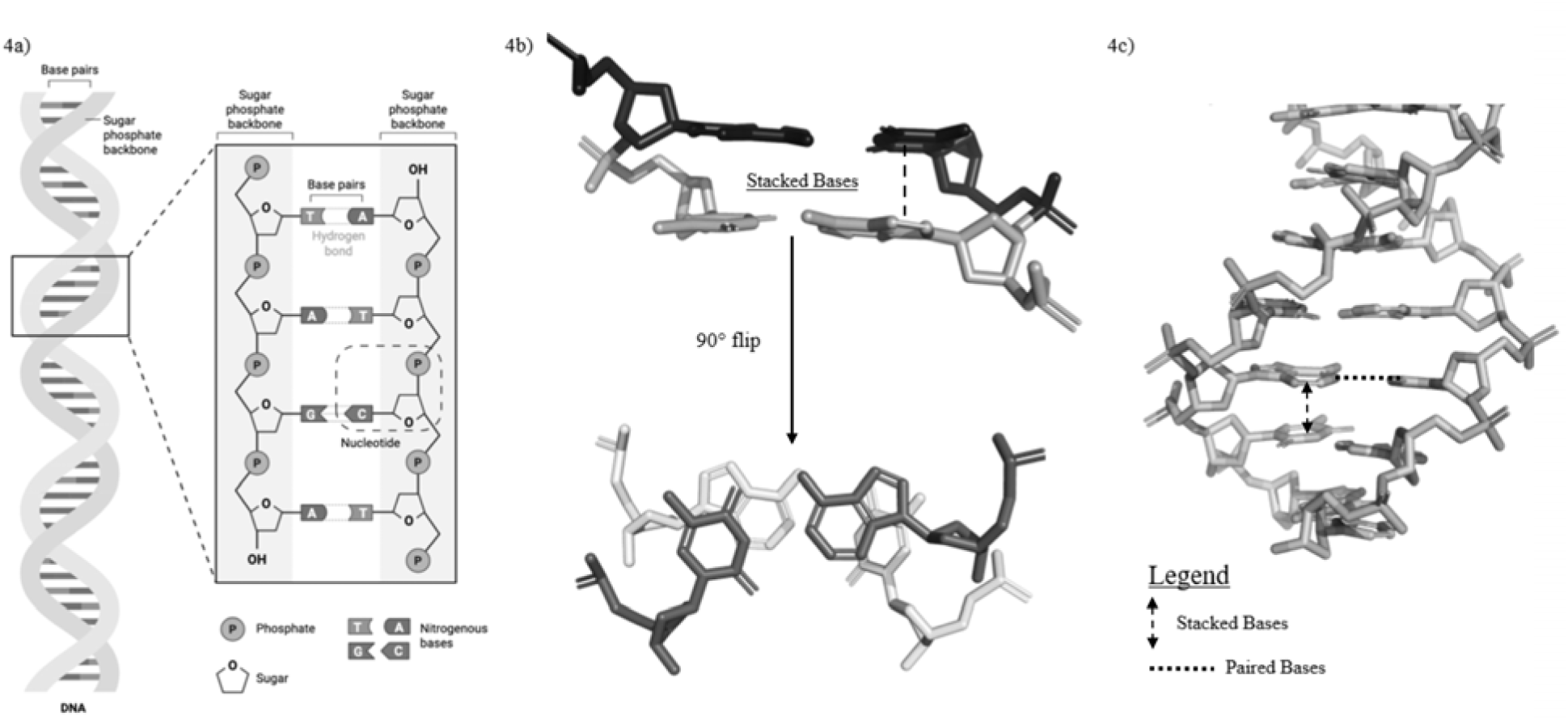
Representation of figure biases in illustrating the forces that stabilize double stranded DNA. a) Representation of a figure clearly indicating base pairing, taken from BioRender. 4b) Representation of a figure clearly indicating base stacking from a side-on and birds’-eye view. 4c) Representation of a figure clearly indicating base stacking with dashed double headed arrows and pairing with a dotted line, as indicated by the legend. dsDNA was generated in PyMOL using the “fnab” command.

Three coders (two students and the instructor) independently re-categorized the 35 textbooks based on the decision tree into biased towards base pairing, base stacking, or unbiased in both text and figure. Coders met in person and cross examined each other’s results. For any discrepancies in categorization, the coders debated by presenting evidence to support their decision until unanimous decisions were reached, which eventually formed part of the result section of this paper.

### Literature Review

Literature review was conducted in stages as part of the scaffolded writing assignments for the year-long laboratory experience as described before. The goal for literature search and review was for students to develop a more holistic and nuanced view on DNA stability, rather than on the comprehensiveness of the literature review.

Students working in small research teams were instructed to search for key words: such as base pairing, base stacking, DNA stability on PudMed and Google Scholar, prioritizing publications from the last ten years. Students were provided with freedom to generate their own bibliography and initial draft of literature review. During revision of their final manuscript, the bibliographies from different student groups were pooled and shared with all students to expand the scope of their literature review. Three themes on DNA structure and function, environmental effects on DNA, and DNA-protein interaction emerged from students’ literature reviews were presented in this paper.

## Results & Discussion

### 1. How might textbook representations both in writing and in figure contribute to students’ oversimplified view on DNA stability?

Through a systematic analysis of both textbook content and figures, 35 textbooks were allocated into five out of the nine categories as outlined in the decision tree and summarized in Table 1. This initial analysis revealed that the largest proportion of textbooks explicitly favoured base pairing both in text and in figure as the dominant force that stabilizes the dsDNA, which might have influenced instructors and thus predisposed students to favour base paring as the dominant force that stabilizes dsDNA during teaching and learning respectively.

**Table 1.** Textbooks sorted into categories based on the decision tree described in the Methods.

| Category | Number of Textbooks |
| --- | --- |
| Texts favour base stacking; Figures favour base pairing | 5 |
| Texts favour base stacking; Figures unbiased | 5 |
| Texts unbiased; Figures favour base pairing | 6 |
| Text unbiased; Figures unbiased | 7 |
| Texts favour base pairing; Figures favour base pairing | 12 |

We further analyzed the texts and figures separately to reveal additional trends that might be masked from the previous overall analysis. When looking only at the texts in the 35 textbooks analyzed, we found a relatively equal proportion of books favouring pairing, stacking, and unbiased descriptions of dsDNA (Figure 5). This distribution represented that in two thirds of the textbooks, the authors showed a clear preference of one force over the other. Notably, there were striking contradictions across textbooks which could cause confusion among students. For instance, Elliott & Elliott (2005) assert unequivocally, “DNA almost always exists as a double strand … What holds them together? The answer is complementary base pairing.” In stark contrast, Pratt & Cornely (2021) state, “Instead, stability depends mostly on stacking interactions, which are a form of van der Waals interactions, between adjacent base pairs.”

**Figure 5.**
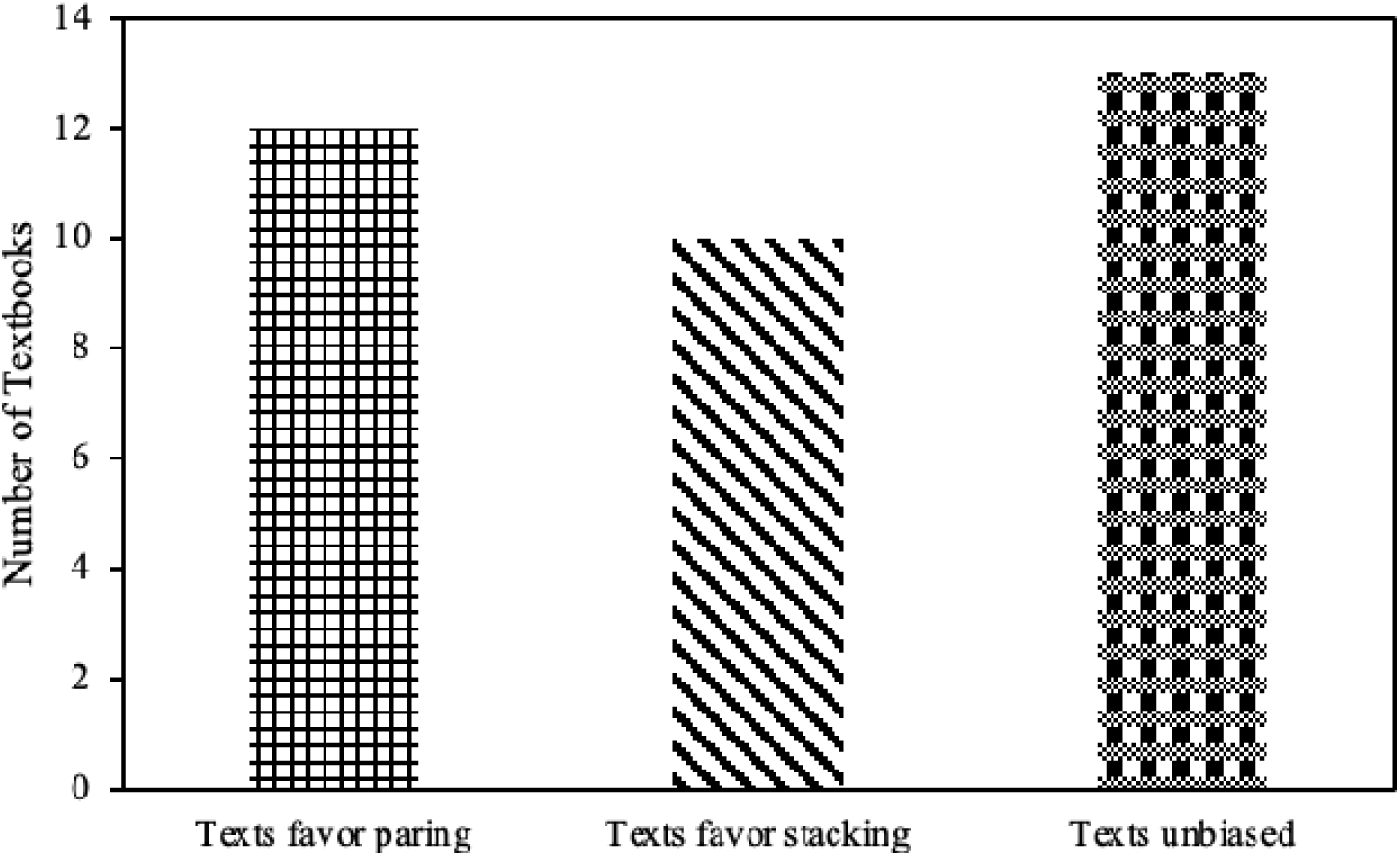
Number of textbooks that were assigned as having written texts biased towards pairing, stacking, or having no bias.

A closer look at textbooks reveals a clear divide. Voet & Voet (2011) emphasize the contribution of H-bonds, noting that the melting point for double stranded DNA “…increases linearly with the mole fraction of G-C base pairs, which indicates that triply hydrogen bonded G-C base pairs are more stable than A-T base pairs.” Conversely, Miesfeld & McEvoy (2021) challenges this view by stating “Sequences rich in G-C base pairs have more stability than do sequences rich in A-T base pairs primarily due to more favourable base stacking interactions (not because of differences in the number of hydrogen bonds).” Similarly, Devlin (2002) acknowledges the historical neglect of stacking, stating “The relative importance of base stacking versus hydrogen bonding in stabilizing the double helix was not always appreciated … experiments with reagents that reduce the stability of the double helix illustrate the greater importance of stacking interactions.” When these claims are presented across different sources without reconciliation, they might contribute to a conflicting educational narrative.

When analysing only the figures from the 35 textbooks, we found two thirds of the textbooks’ figures favoured base pairing, while the remaining third showed a more unbiased representation of both base pairing and stacking (Figure 6). Taken together with the text-only analysis, we found that even though 23 textbooks in writing favoured either base stacking or unbiased—for which we would expect their figures to support either base stacking or an unbiased illustration, respectively—in practice only 12 of those textbooks showed unbiased figure representation, while no book showed figures that favoured base stacking. These results represent a nearly 50% reduction in highlighting the contribution of base stacking in stabilizing dsDNA. This discrepancy illuminates a critical issue—visual aids disproportionately favour base pairing, thereby potentially creating a disconnect between textual explanations and visual illustrations during learning.

**Figure 6.**
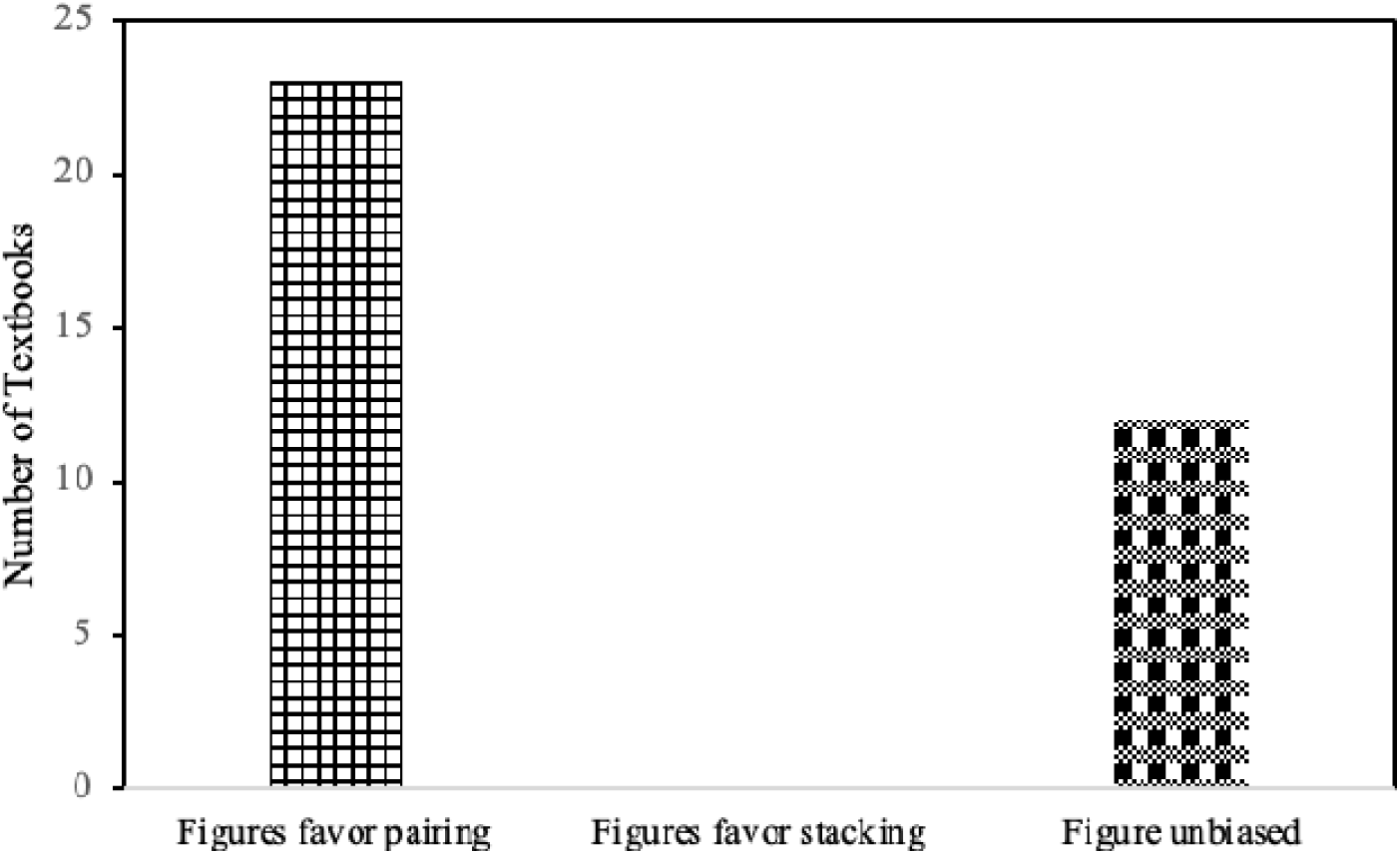
Number of textbooks that were assigned as having figures biased towards pairing, stacking, or having no bias.

It can be concluded that the disparity between textbook content and figures aligns with the students’ preference for base pairing as the dominant force stabilizing dsDNA. This suggests that the treatment of DNA stability in textbooks might influence student understanding. Textbook figures, often powerful instructional tools for reinforcing concepts, overwhelmingly represent base pairing as the main DNA stabilizing force. As visual learning strongly influences cognitive retention, students and instructors may be unintentionally guided towards emphasizing base pairing, despite the texts recognizing stacking forces as an equivalent if not more stabilizing force. Thus, the prevalence of base-pair-focused figures appears to play a critical role in shaping student understanding and their subsequent misconceptions about DNA stabilization. Addressing this imbalance through promoting clearer, more balanced visual representations in educational materials could help rectify student misunderstandings, ensuring a more complete comprehension of the true nature of DNA stability.

Our textbook analysis described above also has limitations. First, to simplify our analysis, we have limited our analysis to the introductory chapter on DNA structure and function. This means we did not include further chapters and discussion on the molecular mechanisms revolving around central dogma, such as DNA replication and transcription, where both base paring and base stacking clearly play important roles. As a result, any holistic discussion of DNA stability in later chapters would not have been captured in our analysis. Second, our analysis is based on the available access to physical and online textbooks. In particular, for physical textbooks from a variety of different publishers, we do not always have the newest edition. As a result, the current analysis may not be reflective of the most up to date representation on DNA stability in textbooks. Finally, we excluded figure captions from our analysis, because many figure captions would mention base stacking but not highlight it in the actual figure. This decision may have also eliminated the possibility that students could benefit from reading the captions even without a clear label in the figure, meaning reading the involvement of base stacking in the figure captions is enough to take a note of its important roles in stabilizing DNA.

### 2. How does the current literature represent the intermolecular forces involved in DNA stability?

To help students and teachers to approach the concept of DNA stability from a more holistic standpoint, we have chosen three commonly discussed topics in life science programs to illustrate the interplay of the intermolecular interactions involved in stabilizing the DNA double helical structure, emphasizing base pairing and stacking.

#### Example 1: The Structure-Function Relationship of B-DNA

The overall literature review suggests there is a strong interplay between the intermolecular forces stabilizing DNA, highlighting that both base pairing and stacking are necessary for the stability and functionality of DNA (Zhou et al., 1999; Guckian et al., 2000). Matray et al. utilized non-H-bonding analogs *in vitro* and observed that base stacking interactions alone could only stabilize for the loss of 40% of H-bonding, after which the DNA adopted a new structure, referred to as a hairpin (1998). In fact, it has been found that base stacking can enhance H-bond strength, suggesting that the two forces are mutually beneficial (Mignon et al., 2005).

However, it should be acknowledged that the previous claims do not reflect B-DNA in biological systems, and this may be the source of discrepancy amongst findings in research reporting on DNA stability. It is not unlikely that in controlling experimental conditions, contributions of one force have been overlooked or influenced by experimental parameters. This can be addressed by considering the function of B-DNA as genetic material and how this is influenced by structure, sequence, and environmental conditions.

In a cellular environment, DNA has been observed in multiple structural formations, including A, Z, and B-form DNA. Unlike A– and Z-DNA, B-DNA is favoured in aqueous environments. This is attributed to the lack of slide, which can be thought of as a parameter describing the relative deformation of neighbouring base pairs from a completely overlapped orientation (Hunter, 1997). Thus, the lack of slide accommodates hydrophobic interactions between stacked bases and solvation of the major and minor grooves (Hunter, 1993). Despite this, studies have shown that the absence of slide in B-DNA creates unfavourable electrostatic repulsion in a GC base step. This repulsion is not observed in A-DNA, which contains a negative slide, yet there is only a minimal amount of A-DNA observed in the cell (Calladine & Drew, 1984). The abundance of B-DNA proves that the enhanced stacking and solvation effects of B-DNA are more favourable within the cell than simply minimizing electrostatic repulsion. However, this preference for B-DNA *in vivo* does not only arise from base stacking. A study by Kool and colleagues highlights that when replacing up to 40% of the nucleobases in B-DNA with nonpolar analogs incapable of H-bonding, similar stability is achieved, but the degree of solubility is lost (Hirao et al., 2012; Kool et al., 2000). This is because in the absence of H-bonds, non-polar base analogues can achieve a larger extent of overlap, resulting in a lower solvent-accessible surface area (Hirao et al., 2012; Kool et al., 2000; Loakes et al., 1997). Therefore, despite stacking interactions contributing extensively to the stabilization of B-DNA, base pairing is essential for achieving a structure with optimal solubility in a biological context (Kool, 2001; Kool et al.; 2000, Guckian et al., 2000).

The fate of a given stretch of DNA is also contingent on nucleobase composition and hence, relative stability. Stabilization by base stacking and base pairing differs amongst base pairs, resulting in DNA sequences with segregated nucleotide motifs of distinct properties (Jabbari et al., 2019). Specifically, GC-rich sequences are more stable than AT-rich sequences. This is in part due to the additional H-bond in GC pairs, and due to the reinforcement of stacking forces by the overlap of electropositive amide nitrogen and electronegative carbonyl oxygen between diagonal bases in a GC rich sequence (Nieuwland et al., 2022). Furthermore, the presence of both stacking and pairing in B-DNA is crucial for optimizing cellular processes. Considering DNA breathing events that take place during replication and transcription, an AT-rich consensus sequence upstream of the gene has relatively weak stability, enabling the actions of helicase (Gutiérrez-Flores et al., 2017; Krueger et al., 2006). Conversely, sequences rich in GC have enhanced stability, and an extreme example of this is the G rich region in telomere, which forms G-quadruplexes to stabilize and protect the chromosomal ends via enhanced base paring and stacking.

In summary, both base pairing and base stacking are essential for DNA stability, and neither force alone can reinforce the unique structure of DNA. While stacking can compensate for some loss of hydrogen bonding, the extent of stacking compensation is limited by both solubility and structural constrains. Having more base stacking interactions could theoretically stabilize DNA; however, the increased number of hydrophobic nucleobases drastically decrease DNA solubility in water, collapsing the double helical structure. Research further demonstrates the superior stabilizing effect of GC-rich regions in DNA results from both the additional H-bond and increased base stacking. Overall, DNA’s behavior in cells clearly depends on the combined contributions of both base pairing and base stacking, which work together to support DNA structure and biological function.

#### Example 2: Environmental Effect on DNA Stability

Base pairing and base stacking interactions can be influenced by environmental factors, with solvation emerging as a significant area of interest. The impacts of solvation on base pairing have been extensively studied using both computational and experimental methodologies.

Quantum chemical computational studies have allowed researchers to delve into the behaviour of WC base pairs while considering structural and environmental factors. Computational studies of DNA stability indicate that solvation has a destabilizing effect on the formation of DNA duplexes, due to a slight decrease in cavitation energy, which reflects the energy required to create a void in the solvent for the solute. The solvent concentration and type significantly influence base pairing by not only reducing H-bonding but also equilibrating the interactions between H-bonding and π-π stacking for both AT and GC base pairs (Poater et al., 2014; Nieuwland et al., 2022).

Experimental investigations utilizing calorimetry and spectroscopy techniques have supported computational predictions regarding the stability of base pairing. These experiments have shown that the hydration of DNA minor grooves plays a crucial role in stabilizing H-bonds between base pairs (Privalov & Crane-Robinson, 2018). The release of structured water molecules during DNA melting significantly contributes to the enthalpic and entropic aspects of dsDNA stability, underscoring the importance of solvent mediated interactions in base pairing (Privalov & Crane-Robinson, 2018). Additionally, similar studies have provided direct measurements of the strength of DNA H-bonds, confirming that variations in ionic strength within solvent conditions directly affect base pairing (Zhang et al., 2015). Under low-salt conditions, DNA H-bonds demonstrate increased instability, while higher concentrations of NaCl enhance the force required to dissociate complementary strands.

Molecular dynamic simulations of DNA stability have revealed that solvation effects also significantly influence stacking interactions in B-DNA (Poater et al., 2014; Tran et al., 2022). The findings indicate that base stacking interactions are more robust in hydrophobic environments, whereas they weaken in aqueous solvation conditions due to enhanced hydration-induced electrostatic shielding. Free energy calculations of stacked DNA duplexes under varying solvent conditions have confirmed this through demonstrating that water molecules disrupt base stacking by altering the electrostatic potential of π–π interactions (Punnoose et al., 2025).

Single-molecule force spectroscopy experiments provide direct mechanical insights into base-stacking interactions (Zhang et al., 2015). These studies indicate that the forces required to disrupt base stacking in individual DNA molecules depend on solvent conditions and ionic strength. Notably, in high-salt environments, stacking interactions become increasingly strong, while in low-salt conditions, reduced electrostatic shielding results in weakened π–π interactions, leading to enhanced DNA flexibility and instability (Yakovchuk, 2006).

Interestingly, the identity of the salt in the solvent also influences the relative strength of base pairing and stacking in stabilizing B-DNA. Huguet et al. conclude that the presence of the monovalent cation sodium, increases the contribution of stacking interactions towards duplex stability, and the presence of the divalent cation magnesium increased the contribution of base pairing (Bizarro et al., 2012; Huguet et al., 2017; McFail-Isom et al., 1998; Punnoose et al., 2025; Viader-Godoy et al., 2024;). These results suggest that the ionic strength and stability of the duplex’s environment influence the relative strength of its pairing and stacking interactions. This phenomenon is caused by the inhibition of base stacking in the presence of magnesium rather than the activation of base pairing in sodium (Punnoose et al., 2025; Bizarro et al., 2012; Viader-Godoy et al., 2024; McFail-Isom et al., 1998). Magnesium ions in the major groove of B-DNA have been demonstrated to be able to pull nucleobases partially out of the duplex, causing unstacking, which likely plays a key mechanistic role in DNA bending, strand separation, and DNA-protein recognition and complexation (Punnoose et al., 2025; Bizarro et al., 2012; Viader-Godoy et al., 2024; McFail-Isom et al., 1998; Naskar et al., 2019).

Taken results from both computation and experimentation together, we may conclude:

- Water generally weakens both base pairing and base stacking interactions, but with different mechanisms. Water competes with nucleobases for H-bonds; while water hydrates π surfaces and shields electrostatic interactions.
- Salt in solvents generally improves both base pairing and base stacking interaction by reducing the backbone charge repulsion, allowing the two strands to pack more tightly through both base pairing and base stacking.
- Despite of the relative consistency on the solvent effect, recent studies have revealed subtle differences on how ion identity (Na⁺ vs. Mg²⁺) shifts the base pairing and base stacking balance, with Na⁺ enhancing stacking’s contribution to stability and Mg²⁺ enhancing pairing’s contribution.

The discussion on how environmental factors affect DNA stability reinforces that idea that DNA stability arises from a finely tuned balance between base pairing (hydrogen bonds) and base stacking (π–π interactions), where both forces are strongly modulated by the solvent environment and ions.

#### Example 3: DNA Stability Through the Lens of Protein Stability and its Interaction

Protein folding is taught often repeatedly through biochemistry classes, where students understand the fundamental roles of inter and intramolecular forces stabilizing proteins. Applying the understanding of protein stability to DNA stability may be a conducive channel to help students develop a holistic view on how B-DNA is stabilized.

Protein primary structure is linked through peptide (amide) bonds via condensation reactions. Localized amino acids on the linear chain begin to interact with each other to form secondary structures, -helices and β-strands, through H-bonds. Eventually these distal folded motifs and domains coalesce to form a three-dimensional structure driven by the hydrophobic effect, whereby the polar and charged amino acids are exposed to the solvent, maximizing the interactions with water, while hydrophobic amino acids are buried inside, maximizing van der Waals interactions. This is the tertiary structure and often the final structure of a protein. If a protein requires multiple subunits to function, protomers could associate through H-bonds, ionic interactions, disulfide bonds, and van der Waals interactions, forming a quaternary structure. Likewise, in DNA, each polynucleotide is linked through a series of condensation reactions, forming phosphodiester bonds. The two polynucleotides form the well-known double stranded helix due to the hydrophobic effect, reminiscent of protein folding process.

These complementary interactions in stabilizing protein and DNA also dictate how the two molecules interact with each other. For example, DNA helicase directly binds to DNA through both van der Waals interaction and ionic interactions. DNA helicase uses the conserved aromatic amino acids (e.g. Trp or Phe) to establish π-π interactions with the nucleobases and uses electropositive amino acids (e.g. Arg) to neutralize the negatively charged phosphodiester backbone respectively (Bhattacharyya & Keck, 2014). Indeed, the molecular mechanism by which helicases unwind DNA illuminates the importance of both base pairing and base stacking in maintaining DNA stability.

In both prokaryotes and eukaryotes, the active form of DNA helicase is a hexamer that encircles a single strand of DNA completely (Miesfeld & McEvoy, 2021). Helicase binds to only one of the DNA strands and moves in a 5’-3’ direction. The DnaB helicase in E. coli functions by forming a barrel-like structure through which the nucleic acid passes. ATP provides the energy for translocation along the DNA, with the ATP binding pocket found at the interfaces between the monomers. To separate the two strands of DNA, a helicase must break the hydrogen bonds holding the helix together (Patel & Donmez, 2006). In doing so, it must associate with the single-stranded DNA passing through the helicase to prevent the helix from re-forming. In the case of DnaB, 12 nucleotides of the single-stranded DNA are associated with the helicase at any given time, with each of the six monomers bound to two nucleotides (Miesfeld & McEvoy, 2021).

While helicases are traditionally recognized for their role in disrupting base pairing, their activity has equally significant effects on base stacking—the other essential contributor to DNA stability. The mechanical act of unwinding generates torsional strain, supercoiling, and local distortion within the DNA duplex (Duguet, 1997). These conformational stresses force base stacking out of their optimal coplanar arrangement, weakening π–π interactions and destabilizing stacking. Additionally, helicases create stretches of ssDNA during unwinding, and ssDNA inherently exhibits diminished stacking because bases lack the geometric constraints imposed by Watson–Crick pairing (Brosh, 2015). Thus, helicase progression induces a cascading loss of both stabilizing forces: hydrogen bonds are broken directly, and stacking is weakened both mechanically and structurally.

Taken together, helicase activity provides a compelling biological demonstration of the central argument advanced in this project: DNA stability does not arise from base pairing or base stacking alone but from the interdependent contributions of both. Helicases must overcome hydrogen bonding to initiate strand separation, yet efficient unwinding also depends on disrupting stacking interactions that resist conformational deformation. When helicases destabilize pairing, stacking becomes more vulnerable; when stacking is disrupted, fewer hydrogen bonds need to be broken to propagate strand separation. Neither interaction alone can account for the overall resistance of duplex DNA to unwinding; rather, helicases succeed because they simultaneously weaken both stabilizing forces.

In summary, the mechanism of DNA helicases underscores the inseparability of base pairing and base stacking in maintaining the structural integrity of the double helix. Their coordinated actions reveal that DNA stability is a composite property arising from the interplay between these interactions—precisely the relationship demonstrated by computational, thermodynamic, and singlelllmolecule studies described earlier.

## Conclusion

Through our investigation of student conceptual bias regarding the role of base pairing in stabilizing doublelllstranded DNA, we identified that a major source of misunderstanding arises from the oversimplified way DNA is presented in many educational materials. Textbook figures frequently highlight hydrogen bonding between complementary bases while neglecting the equally essential contribution of base stacking. This visual emphasis might lead students to overestimate the role of hydrogen bonding and underestimate the stabilizing forces provided by π–π interactions.

To address this gap, students examined three foundational topics commonly taught in the life sciences—DNA structure–function relationships, environmental factors affecting DNA stability, and helicaselllmediated DNA unwinding. Across each topic, we demonstrated that base pairing and base stacking operate not as isolated forces but as deeply intertwined contributors to the physical and functional stability of DNA. This integrated perspective reveals that DNA behaves as a dynamic system whose stability reflects the balance of multiple interactions influenced by hydration, ionic conditions, sequence composition, and mechanical forces. By adopting a holistic approach, our discussion underscores that base pairing and base stacking are both crucial for ensuring DNA’s structural integrity, proper solubility in the cellular environment, and responsiveness to biological processes such as replication and transcription.

## Glossary

**X-ray diffraction (XRD)** is a method for visualizing how atoms are arranged in a solid (like a crystal) by shining X-rays at it and measuring the pattern of spots the X-rays make after they scatter. The spot pattern can be used to reconstruct the 3D structure of the model.

A **phosphodiester bond (or linkage)** is the strong covalent bond that connects one nucleotide to the next in DNA. It forms when a phosphate group bridges the sugar of one nucleotide to the sugar of the next, creating a repeating sugar-phosphate chain.

**Major and minor grooves** are the two unequal grooves that spiral along the outside of the DNA double helix. Because the two strands are not positioned symmetrically, the major groove is wider and the minor groove is narrower, proteins often use these grooves to read DNA sequences and bind.

**Quantum chemical computational studies** are calculations that predict how atoms and electrons behave using the rules of quantum mechanics. They are used to estimate things like bond strengths, charge distributions, and how strongly molecules attract or repel each other at the electron level.

**Calorimetry** is a technique that measures heat absorbed or released during a process, such as DNA melting or a protein binding to DNA. By tracking heat changes, calorimetry reveals how energetically favourable an interaction is and can separate contributions like **enthalpy** (heat) and **entropy** (disorder).

**Spectroscopy** is a set of methods that probe molecules by observing how they interact with light (or other forms of radiation), such as absorbing, emitting, or scattering it. The resulting signals act like molecular fingerprints that report on structure, composition, and changes over time.

**Intermolecular forces** are noncovalent attractions (or repulsions) between separate molecules or different parts of large molecules. Common intermolecular forces include:

- **Hydrogen bonds** are directional attractions where a hydrogen attached to a nitrogen, oxygen, or fluorine atom is attracted to another electronegative atom.
- **Ionic interactions** are attractions between fully charged groups (positive and negative).
- **Dipole-Dipole** interactions are attractions between polar molecules with partial charges.
- **π–π interactions** are attractive, noncovalent forces between the aromatic ring systems of nucleobases (A, T, G, C). These interactions occur when the πlllelectron clouds of stacked bases overlap.

**Van der Waals (vdW) Interactions** are weak short-range attractive forces between atoms that come from shifts in their electron clouds. Individually they are small, but many van der Waals contacts together can strongly stabilize how molecules pack and stick together. VdW interactions are also known by the term London Dispersion Forces, in this paper this term is not used.

**Molecular dynamics (MD) simulations** are computer run time-dependant models of molecular motions that tracks how atoms move over time under physical forces. MD is used to study flexibility, folding, binding, and how molecules respond to temperature, salt, water, or mechanical stress.

**Cavitation energy** is the energy required to form a cavity (empty space) in a solvent so that a solute, such as a DNA base or base pair, can be accommodated.

A **hydration shell** is a layer of water molecules that surrounds the DNA duplex. These water molecules interact with the DNA backbone and grooves through hydrogen bonding.

**Slide** measures the sideways displacement of one base pair relative to the next, along an axis perpendicular to the helix. It indicates how much bases shift sideways instead of sitting perfectly aligned.

**Twist** is the rotation between adjacent base pairs around the helical axis, essentially describing how tightly DNA is wound.

**Roll** measures the bending of one base pair relative to the next, like opening a book at the spine. It describes curvature toward the major or minor groove.

## Works Cited

1. Ahern-, K., Rajagopal, I., & Tan, T. (2018). Biochemistry: Free for all. Open Textbook Library. https://open.umn.edu/opentextbooks/textbooks/biochemistry-free-for-all-ahern

2. Alberts, B., Heald, R., Johnson, A., Morgan, D., Raff, M., Rogers, K., Peter, W., Wilson, J., & Hunt, T. (2022). Molecular biology of the cell (7th ed.). W. W. Norton & Company.

3. Anthony-Cahill, S. J., Appling, D. R., & Mathews, C. K. (2019). Biochemistry: Concepts & connections (2nd ed.). Pearson.

4. Armstrong, F. B. (1983). Biochemistry (2nd ed.). Oxford University Press.

5. Astbury, W. T., & Bell, F. O. (1938). X-ray studies of thymonucleic acid. Nature, 141, 747–748. 10.1038/141747b0

6. Berg, J. M., Gatto, G. J., Jr., Hines, J. K., Tymoczko, J. L., & Stryer, L. (2023). Biochemistry (10th ed.). Macmillan Learning.

7. Berg, J. M., Tymoczko, J. L., & Stryer, L. (2002). Biochemistry (5th ed.). W. H. Freeman and Company.

8. Bhattacharyya, B., & Keck, J. (2014). Grip it and rip it: Structural mechanisms of DNA helicase substrate binding and unwinding. Protein Science, 23(11), 1498–1507. 10.1002/pro.2533

9. Bizarro, C. V., Alemany, A., & Ritort, F. (2012). Non-specific binding of Na? and Mg²? to RNA determined by force spectroscopy methods. Nucleic Acids Research, 40(14), 6922–6935. 10.1093/nar/gks289

10. Boyer, R. (1999). Concepts in biochemistry. Thomson-Brooks/Cole.

11. Brosh, R. M., Jr. (2015). DNA helicases involved in DNA repair and their roles in cancer. Nature Reviews Cancer, 13(8), 542–558. 10.1038/nrc3560

12. Calladine, C., & Drew, H. (1984). A base-centred explanation of the B-to-A transition in DNA. Journal of Molecular Biology, 178(3), 773–782. 10.1016/0022-2836(84)90251-1

13. Campbell, M. K., & Farrell, S. O. (2003). Biochemistry (4th ed.). Thomson-Brooks/Cole.

14. Campbell, N. A., & Reece, J. B. (2002). Biology (6th ed.). Pearson.

15. Cox, M. M., Hoskins, A. A., Viel, A., & Simcox, J. (2025). Lehninger biochemistry: Core concepts & applications. Macmillan Learning.

16. Cox, M.M., Doudna, J.A., O’Donnell, M. (2015). Molecular biology: Principles and practices (2nd ed.). Macmillan Learning.

17. Crane-Robinson, C. (2022). Role of water in defining the structure and properties of B-form DNA. Crystals, 12(6), 818. 10.3390/cryst12060818

18. Dahm, R. (2005). Friedrich Miescher and the discovery of DNA. Developmental Biology, 278(2), 274–288. 10.1016/j.ydbio.2004.11.028

19. Devlin, T. M. (Ed.). (2002). Textbook of biochemistry with clinical correlations (5th ed.). Wiley-Liss.

20. Dow, J., Lindsay, G., & Morrison, J. (1996). Biochemistry: Molecules, cells and the body. Pearson.

21. Duguet, M. (1997). When helicase and topoisomerase meet! Journal of Cell Science, 110(12), 1345–1350. 10.1242/jcs.110.12.1345

22. Elliott, W. H., & Elliott, D. C. (1997). Biochemistry and molecular biology (1st ed.). Oxford University Press.

23. Elliott, W. H., & Elliott, D. C. (2005). Biochemistry and molecular biology (3rd ed.). Oxford University Press.

24. Ferrier, D. R. (2014). Biochemistry (6th ed.). Wolters Kluwer Health/Lippincott Williams & Wilkins.

25. Feughelman, M., Langridge, R., Seeds, W. E., Stokes, A. R., Wilson, H. R., Hooper, C. W., Wilkins, M. H. F., Barclay, R. K., & Hamilton, L. D. (1955). Molecular structure of deoxyribose nucleic acid and nucleoprotein. Nature, 175(4463), 834–838. 10.1038/175834a0

26. Franklin, R. E., & Gosling, R. G. (1953). Molecular configuration in sodium thymonucleate. Nature, 171, 740–741. 10.1038/171740a0

27. Friedman, R. A., & Honig, B. (1992). The electrostatic contribution to DNA base-stacking interactions. Biopolymers, 32(2), 145–159. 10.1002/bip.360320205

28. Garrett, R. H., & Grisham, C. M. (2005). Biochemistry (3rd ed.). Cengage Learning.

29. Guckian, K. M., Schweitzer, B. A., Ren, R. X., Sheils, C. J., Tahmassebi, D. C., & Kool, E. T. (2000). Factors contributing to aromatic stacking in water: Evaluation in the context of DNA. Journal of the American Chemical Society, 122(10), 2213–2222. 10.1021/ja9934854

30. Gutiérrez-Flores, J., Hernández-Lemus, E., Cortés-Guzmán, F., & Ramos, E. (2020). Do weak interactions affect the biological behavior of DNA? A DFT study of CpG island–like chains. Journal of Molecular Modeling, 26(10). 10.1007/s00894-020-04501-6

31. Gutiérrez-Flores, J., Ramos, E., Mendoza, C. I., & Hernández-Lemus, E. (2017). Electronic properties of DNA: Description of weak interactions in TATA-box-like chains. Biophysical Chemistry, 233, 26–35. 10.1016/j.bpc.2017.11.008

32. Harvey, R. A. & Ferrier, D. (2013). Biochemistry (5th ed.). Wolters Kluwer.

33. Hati, S., & Bhattacharyya, S. (2024). Writing a literature review as a class project in an upper-level undergraduate biochemistry course. Biochemistry and Molecular Biology Education, 52(3), 311–316. 10.1002/bmb.21814

34. Hirao, I., Kimoto, M., & Yamashige, R. (2012). Natural versus artificial creation of base pairs in DNA: Origin of nucleobases from the perspectives of unnatural base pair studies. Accounts of Chemical Research, 45(12), 2055–2065. 10.1021/ar200257x

35. Horton, H. R., Moran, L. A., Perry, M. D., & Scrimgeour, K. G. (2012). Principles of biochemistry (5th ed.). Pearson.

36. Huguet, J. M., Ribezzi-Crivellari, M., Bizarro, C., & Ritort, F. (2017). Derivation of nearest-neighbor DNA parameters in magnesium from single molecule experiments. Nucleic Acids Research, 45(22), 12921–12931. 10.1093/nar/gkx1161

37. Hunter, C. A., & Lu, X. (1997). DNA base-stacking interactions: A comparison of theoretical calculations with oligonucleotide X-ray crystal structures. Journal of Molecular Biology, 265(5), 603–619. 10.1006/jmbi.1996.0755

38. Hunter, C. A. (1993). Sequence-dependent DNA structure. Journal of Molecular Biology, 230(3), 1025–1054. 10.1006/jmbi.1993.1217

39. Jabbari, K., Chakraborty, M., & Wiehe, T. (2019). DNA sequence-dependent chromatin architecture and nuclear hubs formation. Scientific Reports, 9(1). 10.1038/s41598-019-51036-9

40. Jakubowski, H., & Flatt, P. (2018, December 2). Fundamentals of biochemistry. Biology LibreTexts. https://bio.libretexts.org/Bookshelves/Biochemistry/Fundamentals_of_Biochemeistry_(Jakubowski_and_Flatt)

41. Klostermeier, D., & Rudolph, M. G. (2018). Biophysical chemistry (1st ed.). CRC Press, Taylor & Francis Group.

42. Kool, E. T., Morales, J. C., & Guckian, K. M. (2000). Mimicking the structure and function of DNA: Insights into DNA stability and replication. Angewandte Chemie International Edition, 39(6), 990–1009. 10.1002/(SICI)1521-3773(20000317)39:6<990::AID-ANIE990>3.0.CO;2-0

43. Kool, E. T. (2001). Hydrogen bonding, base stacking, and steric effects in DNA replication. Annual Review of Biophysics and Biomolecular Structure, 30(1), 1–22. 10.1146/annurev.biophys.30.1.1

44. Krueger, A., Protozanova, E., & Frank-Kamenetskii, M. D. (2006). Sequence-dependent basepair opening in DNA double helix. Biophysical Journal, 90(9), 3091–3099. 10.1529/biophysj.105.078774

45. Li, S., Cooper, V. R., Thonhauser, T., Lundqvist, B. I., & Langreth, D. C. (2009). Stacking interactions and DNA intercalation. Journal of Physical Chemistry, 113(32), 11166–11172. 10.1021/jp905765c

46. Loakes, D., Hill, F., Brown, D. M., & Salisbury, S. A. (1997). Stability and structure of DNA oligonucleotides containing non-specific base analogues. Journal of Molecular Biology, 270(3), 426–435. 10.1006/jmbi.1997.1129

47. Matray, T. J., & Kool, E. T. (1998). Selective and stable DNA base pairing without hydrogen bonds. Journal of the American Chemical Society, 120(24), 6191–6192. 10.1021/ja9803310

48. McCarty, M., & Avery, O. T. (1945). Studies on the chemical nature of the substance inducing transformation of pneumococcal types. Journal of Experimental Medicine, 83(2), 89–96. 10.1084/jem.83.2.89

49. McFail-Isom, L., Shui, X., & Williams, L. (1998). Divalent cations stabilize unstacked conformations of DNA and RNA by interacting with base systems. Biochemistry, 37(49), 17105–17111. 10.1021/bi982201

50. McKee, T., & McKee, J. R. (2016). Biochemistry: The molecular basis of life (6th ed.). Oxford University Press.

51. McMurry, J., Madsen, S., Ballantine, D., Hoeger, C., & Peterson, V. (2013). Fundamentals of general, organic, and biological chemistry (7th ed.). Pearson.

52. McMurry, J., Madsen, S., Ballantine, D., Hoeger, C., & Peterson, V. (2017). Fundamentals of general, organic, and biological chemistry (8th ed.). Pearson.

53. Miesfeld, R. L., & McEvoy, M. M. (2017). Biochemistry (1st ed.). W. W. Norton & Company.

54. Miesfeld, R. L., & McEvoy, M. M. (2021). Biochemistry (2nd ed.). W. W. Norton & Company.

55. Mignon, P., Loverix, S., Steyaert, J., & Geerlings, P. (2005). Influence of the p-interaction on the hydrogen bonding capacity of stacked DNA/RNA bases. Nucleic Acids Research, 33(6), 1779–1789. 10.1093/nar/gki317

56. Naskar, S., Gosika, M., Joshi, H., & Maiti, P. K. (2019). Tuning the stability of DNA nanotubes with salt. Journal of Physical Chemistry C, 123(14), 9461–9470. 10.1021/acs.jpcc.8b10156

57. Nelson, D. L., & Cox, M. M. (2021). Lehninger principles of biochemistry (8th ed.). W. H. Freeman and Company.

58. Nelson, D. L., & Cox, M. M. (2008). Lehninger principles of biochemistry (5th ed.). W. H. Freeman and Company.

59. Nieuwland, C., Hamlin, T. A., Guerra, C. F., Barone, G., & Bickelhaupt, F. M. (2022). B-DNA structure and stability: The role of nucleotide composition and order. ChemistryOpen, 11(2). 10.1002/open.202100231

60. Panini, S. R. (2013). Medical biochemistry: An illustrated review. Thieme Medical Publishers.

61. Patel, S. S., & Donmez, I. (2006). Mechanisms of helicases. Journal of Biological Chemistry, 281(27), 18265–18268. 10.1074/jbc.R600008200

62. Peirce, B. A. (2019). Genetics: A conceptual approach (7th ed.). W. H. Freeman and Company.

63. Petersheim, M., & Turner, D. H. (1983). Base-stacking and base-pairing contributions to helix stability: Thermodynamics of double-helix formation with CCGG, CCGGp, CCGGAp, ACCGGp, CCGGUp, and ACCGGUp. Biochemistry, 22(2), 256–263. 10.1021/bi00271a004

64. Poater, J., Swart, M., Bickelhaupt, F. M., & Guerra, C. F. (2014). B-DNA structure and stability: The role of hydrogen bonding, p–p stacking interactions, twist-angle, and solvation. Organic & Biomolecular Chemistry, 12(26), 4691–4700. 10.1039/c4ob00427b

65. Pratt, C. W., & Cornely, K. (2021). Essential biochemistry (5th ed.). John Wiley & Sons.

66. Privalov, P. L., & Crane-Robinson, C. (2018). Forces maintaining the DNA double helix and its complexes with transcription factors. Progress in Biophysics and Molecular Biology, 135, 30–48. 10.1016/j.pbiomolbio.2018.01.007

67. Punnoose, J. A., Cole, D., Melfi, T., Morya, V., Madhanagopal, B. R., Chen, A. A., Vangaveti, S., Chandrasekaran, A. R., & Halvorsen, K. (2025). Tuning the stability of DNA tetrahedra with base stacking interactions. Nano Letters, 25(9), 3605–3612. 10.1021/acs.nanolett.4c06548

68. Shafee, T. (2019). 3xDNA-DSSR: A resource for structural bioinformatics of nucleic acids. National Institute of Medical Science. https://x3dna.org/highlights/pauling-triplex-model-of-nucleic-acids-is-available-in-3dna

69. Tanford, C. (1978). The hydrophobic effect and the organization of living matter. Science, 200(2), 1012–1018. 10.1126/science.653353

70. Tansey, J. T. (2020). Biochemistry: An integrative approach with expanded topics. John Wiley & Sons, Inc.

71. Timberlake, K. C. (2010). General, organic, and biological chemistry: Structures of life (3rd ed.). Prentice Hall.

72. Tran, B., Cai, Y., Janik, M. J., & Milner, S. T. (2022). Hydrogen bond thermodynamics in aqueous acid solutions: A combined DFT and classical force-field approach. Journal of Physical Chemistry A, 126(40), 7382–7398. 10.1021/acs.jpca.2c04124

73. Tymoczko, J. L., Berg, J. M., Gatto, G. J., Jr., & Stryer, L. (2019). Biochemistry: A short course (4th ed.). W. H. Freeman and Company.

74. Viader-Godoy, X., Manosas, M., & Ritort, F. (2024). Stacking correlation length in single-stranded DNA. Nucleic Acids Research, 53(14), 13243–13254. 10.1093/nar/gkae934

75. Voet, D., & Voet, J. G. (2011). Biochemistry (4th ed.). John Wiley & Sons.

76. Voet, D., Voet, J. G., & Pratt, C. W. (2016). Fundamentals of biochemistry: Life at the molecular level (5th ed.). John Wiley & Sons.

77. Watson, J. D., & Crick, F. H. (1953). Molecular structure of nucleic acids: A structure for deoxyribose nucleic acid. Nature, 171(4356), 737–738. 10.1038/171737a0

78. Yakovchuk, P. (2006). Base-stacking and base-pairing contributions into thermal stability of the DNA double helix. Nucleic Acids Research, 34(2), 564–574. 10.1093/nar/gkj454

79. Zhang, T., Zhang, C., Dong, Z., & Guan, Y. (2015). Determination of base binding strength and base stacking interaction of DNA duplex using atomic force microscope. Scientific Reports, 5(1). 10.1038/srep09143

80. Zhou, H., Yang, Z., & Ou-Yang, Z.-C. (1999). Bending and base-stacking interactions in double-stranded DNA. Physical Review Letters, 82(22), 4560–4563. 10.1103/physrevlett.82.4560

